# Detecting the Indo-Pacific finless porpoises (*Neophocaena phocaenoides*) in Hong Kong waters using environmental DNA

**DOI:** 10.64898/2025.12.21.695310

**Authors:** Xiaoqi Lin, Masayuki Ushio

## Abstract

The Indo-Pacific finless porpoise (*Neophocaena phocaenoides*), a coastal cetacean species widely distributed from the Taiwan Strait to the Persian Gulf, is an iconic species that may represent an overall status indicator of marine ecosystems. However, growing anthropogenic pressures have led to declines in the population of this species in many regions such as Hong Kong waters, highlighting the need for more efficient and reliable monitoring methods. In this study, we applied two environmental DNA (eDNA) analysis methods, species-specific quantitative PCR (qPCR) and cetacean-targeting metabarcoding using µCeta primers, to detect Indo-Pacific finless porpoises’ eDNA in Hong Kong waters. First, we empirically validated the performance of the qPCR primers previously designed in another study. Then, in March 2025, water samples were collected from surface and bottom water layers in 20 locations in the Soko Islands area, a major habitat of the Indo-Pacific finless porpoises. The qPCR analysis detected the finless porpoise eDNA in eight out of 40 samples, but all of the eDNA concentrations were below the limit of quantification. µCeta metabarcoding detected the finless porpoise eDNA from seven out of 40 samples. The qPCR and µCeta metabarcoding approaches showed comparable detection rates for Indo-Pacific finless porpoise. Interestingly, the µCeta metabarcoding method detected a transient cetacean species, the false killer whale (*Pseudorca crassidens*), from a single location. Both methods showed consistent detection rates between surface and bottom water layers, with no statistically significant difference. Overall, the two eDNA analysis methods successfully detected the finless porpoise eDNA in Hong Kong waters, demonstrating the potential of these approaches for monitoring the Indo-Pacific finless porpoises and other cetaceans in this region. Taken together, our findings provide a basis for an eDNA-based cetacean monitoring framework in Hong Kong waters.

## 1. Background

Cetaceans, a group of marine mammals that are fully aquatic throughout their lives, consist of 94 species, including Mysticeti (baleen whales, 15 species) and Odontoceti (toothed whales, dolphins and porpoises: 79 species) [1]. As top predators in the ocean, cetaceans play a crucial role in maintaining the flow of matter and energy and the health of marine ecosystems [2–3]. Thus, changes in the abundance and distribution of cetaceans will alter the structure of marine ecosystems, thereby affecting ecosystem functions such as biogeochemical cycles and carbon sequestration [4–5]. The Indo-Pacific finless porpoise (*Neophocaena phocaenoides*), belonging to the genus *Neophocaena* in the family Phocoenidae, is a common coastal cetacean distributed in shallow waters (<50 m depth) from the Persian Gulf to the Taiwan Strait and southward to Indonesia [6–9]. It typically inhabits tropical to warm temperate coastal environments such as bays, mangrove swamps, estuaries, and large rivers [10].

Hong Kong, located east of the Pearl River Estuary (PRE) and bordering the South China Sea, serves as a key long-term habitat for the Indo-Pacific finless porpoise [11]. Its marine waters span approximately 1,649 km^2^ with a complex coastline of 1,189 km that includes numerous bays, channels, and islands [12]. The region exhibits strong hydrological seasonality: winter is characterized by the northeast monsoon bringing cool, dry air from mainland China, while summer is influenced by the southwest monsoon bringing warm, humid conditions and frequent typhoons [12–13]. Freshwater discharge from the Pearl River creates an estuarine environment in the west, contrasting with the more oceanic conditions in the east, forming an east-west ecological gradient that supports rich biodiversity, including 16 of the 31 marine mammal species recorded in the South China Sea [11,14–15]. Seasonal currents such as the cold Taiwan Current and warm Kuroshio Current further influence the area [12,16]. The Indo-Pacific finless porpoise primarily inhabits the southern and eastern open waters of Hong Kong, while the Indo-Pacific humpback dolphin (*Sousa chinensis*), another resident cetacean in Hong Kong waters, occupies the western waters [10–11,17].

However, due to rapid coastal development and industrialization, the Indo-Pacific finless porpoise faces increasing threats and has been listed as Vulnerable on the IUCN Red List [9,18]. In the PRE region, intense shipping and vessel traffic degrade habitat quality and cause mortality from vessel strikes [6,10], and environmental pollutants, such as DDT and heavy metals, also pose significant risks for the finless porpoises [6]. Local estimates of the finless porpoises in Hong Kong waters indicate a continuous decline from 1996 to 2019 [7], whereas a broader study across the PRE reported an initial decline from 1996 to 2005, followed by a slight recovery from 2006 to 2014 [19–20]. This regional recovery, however, does not explain the concurrent increase in stranding rates in Hong Kong, suggesting that local threats there may be persistent or intensifying [7]. Therefore, more frequent and efficient monitoring is essential to clarify the population status and identify the drivers behind these changes and stranding patterns in Hong Kong waters [7]. However, monitoring finless porpoises is still challenging; their low dorsal ridge and inconspicuous surface behavior make visual surveys, which are weather-dependent and limited to surface-active individuals, difficult and inefficient for consistent detection [6–7, 21–22]. Acoustic monitoring, which can operate continuously under various conditions and detect species-specific vocalizations, offers a promising alternative, but it may be less effective in noisy environments or during periods of low vocal activity [23–24]. Environmental DNA (eDNA) refers to genetic material obtained from environmental samples such as water, and eDNA analysis has emerged as a powerful and efficient tool for biodiversity monitoring [25]. It enables detection of non-invasive species and can even identify intra-specific genetic variations, requiring only minimal field effort such as water collection [26–27]. Also, although careful study design and interpretation are required, eDNA may provide an index for estimating abundance or biomass [28–29]. Given these advantages, applications to cetacean research are growing [30–32]: for instance, eDNA studies have successfully detected deep-diving Cuvier’s beaked whales in surface waters and identified whale subspecies in the open ocean [33–34]. For the Indo-Pacific finless porpoises in Hong Kong waters, two methodological approaches are particularly relevant (Fig. 1): Species-specific qPCR assays offering quantitative analysis of the finless porpoise eDNA [35] and eDNA metabarcoding with cetacean-specific primers (e.g., recently developed µCeta primers) for multi-species detection [36].

**Fig. 1.**
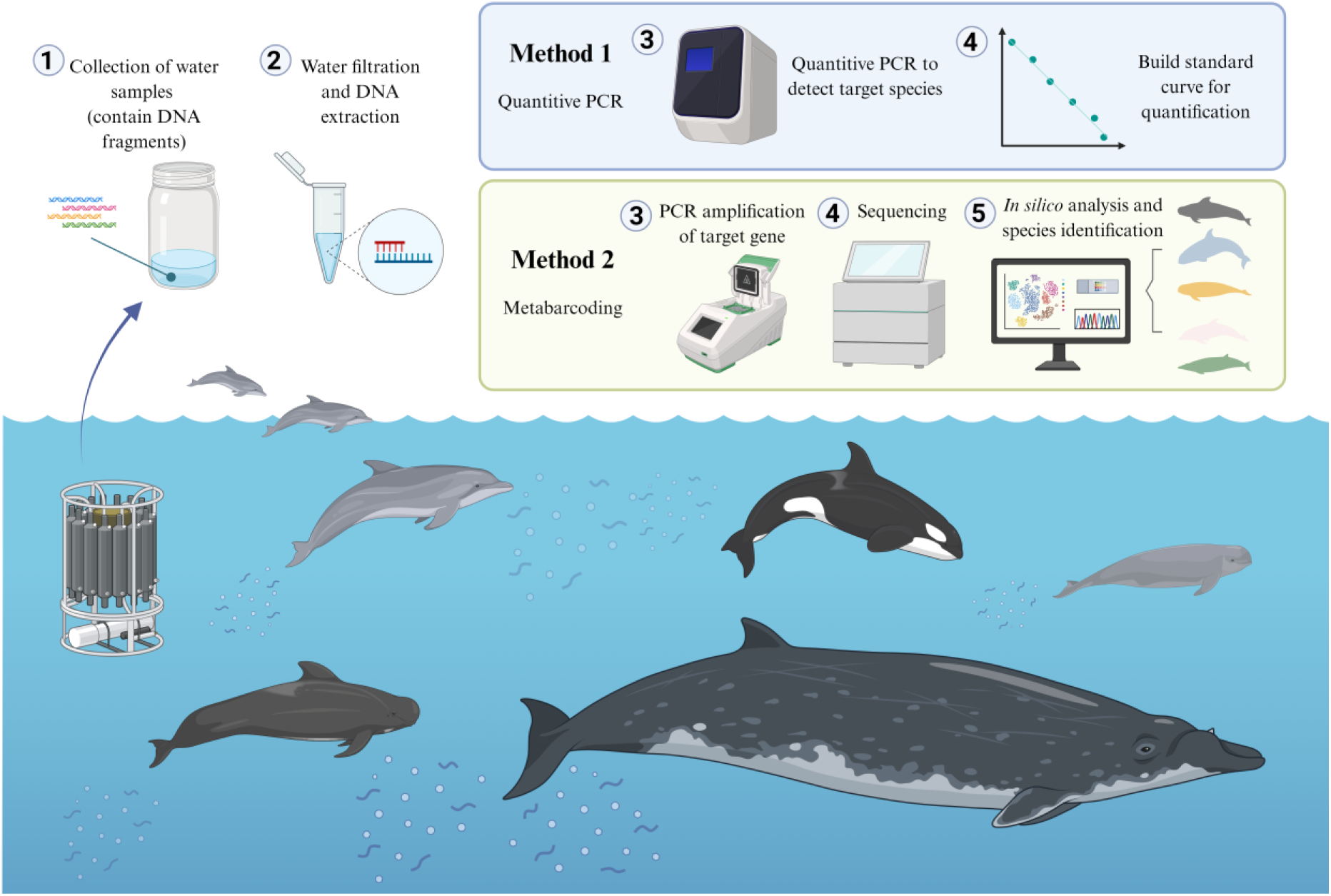
Schematic diagram of eDNA-based cetacean monitoring. A typical procedure is as follows: (1) Collection of water samples from the study area; (2) water filtration and DNA extraction. Then, researchers may utilize either or both of the following methods. Method 1: (3) Quantitative PCR (qPCR) is conducted using a species-specific primer set; (4) standard curve is drawn for eDNA quantification. Method 2: (3) Target genetic region is amplified using a cetacean-specific universal primer set; (4) sequencing of the amplified DNA; (5) sequence analysis and species identification. Created in BioRender. LIN, L. (2025) https://BioRender.com/qe58sir

In this study, we applied two eDNA analysis methods, species-specific qPCR and metabarcoding, to assess the distribution of the Indo-Pacific finless porpoise in Hong Kong waters. qPCR enables the quantification of finless porpoise eDNA, while metabarcoding enables simultaneous detection of multiple cetacean species (e.g., the Indo-Pacific finless porpoise and the Indo-Pacific humpback dolphin). We first evaluated the performance of previously developed finless porpoise-specific qPCR primers. Then, we conducted a field survey around the Soko Islands in Hong Kong waters (which the Indo-Pacific finless porpoises inhabit [37–38]) by collecting surface and bottom water samples at 20 locations. Then, both species-specific qPCR [35] and µCeta metabarcoding [36] were applied to compensate for their respective strengths and weaknesses (i.e., quantitative capacity and multispecies detection), enabling a direct comparison of the sensitivity and reliability of these approaches. Lastly, we examined how detection probability was influenced by water layer (surface vs. bottom) and environmental variables such as salinity and temperature.

## 2. Materials and Methods

### 2.1 Ethics statement

This study used *N. phocaenoides* tissue samples from stranded individuals found in fields to validate species-specific primers for qPCR. All tissue samples were obtained with the necessary permits from the Agriculture, Fisheries and Conservation Department (AFCD) of Hong Kong, Ocean Park Hong Kong (OPHK), and Ocean Park Conservation Foundation Hong Kong (OPCHK). No direct interaction with or sampling from live finless porpoises (or any other cetaceans) occurred during this research. The Soko Islands region is designated as a marine protected area, and we obtained permits for our field survey (i.e., water collection) from AFCD.

### 2.2 qPCR primers and probe and µCeta metabarcoding primers

For qPCR detection, we used finless porpoise-specific qPCR primers and probe designed for the cytochrome b region of the mitochondrial DNA (MT-CYTB, amplicon length = 128 bp; Table 1 and Fig. 2), developed by Hashimoto et al. [35]. The primers were first designed for detecting East Asian finless porpoise (*Neophocaena asiaeorientalis sunameri*), a species closely related to *N*. *phocaenoides*, but we confirmed *in silico* that they can also amplify the Indo-Pacific finless porpoise DNA. Importantly, East Asian finless porpoises are not distributed in Hong Kong waters, and thus, we decided to use the primers and probe to detect Indo-Pacific finless porpoises’ eDNA in Hong Kong waters.

**Table 1.**
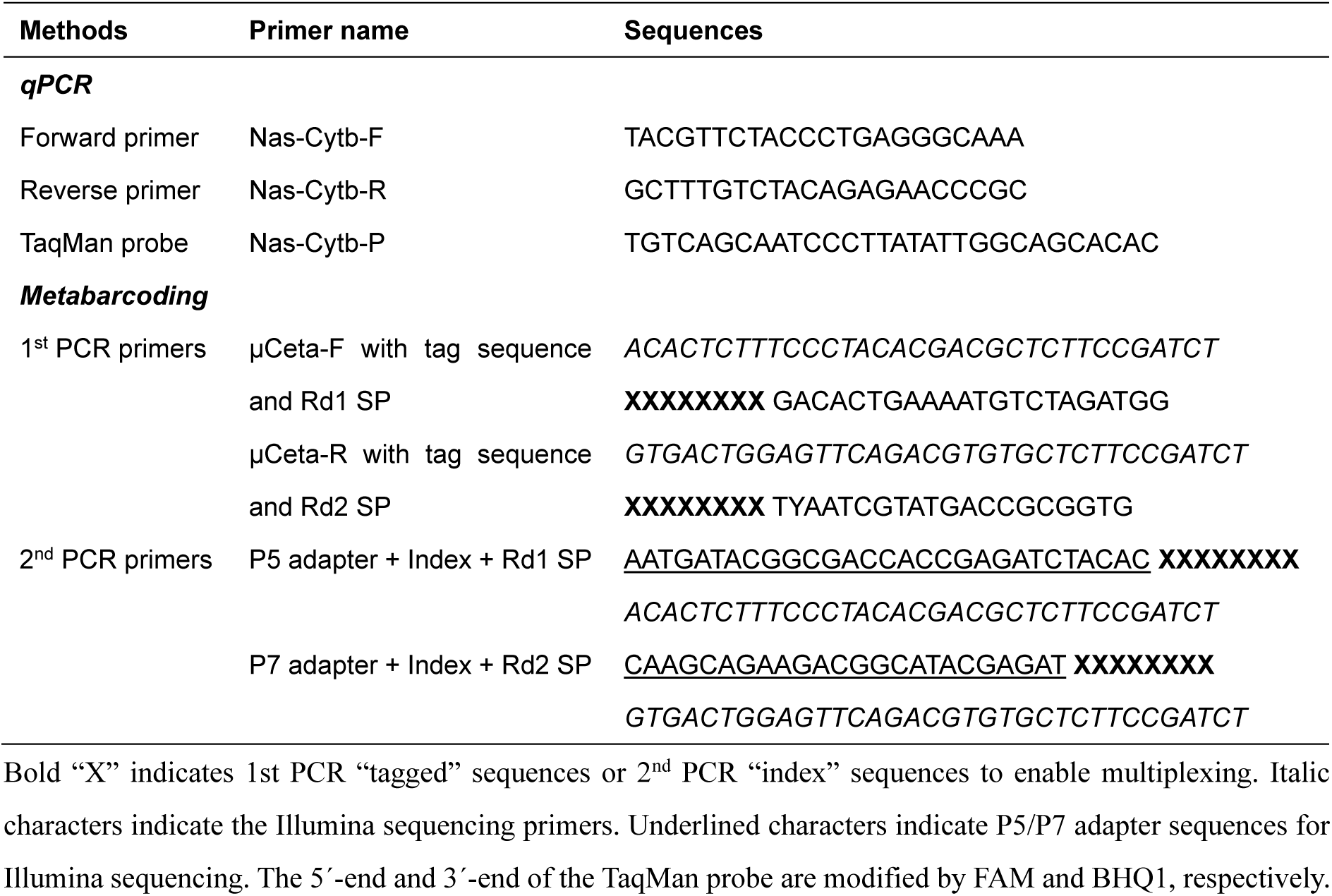
The primers and probe for qPCR and metabarcoding.

**Fig. 2.**
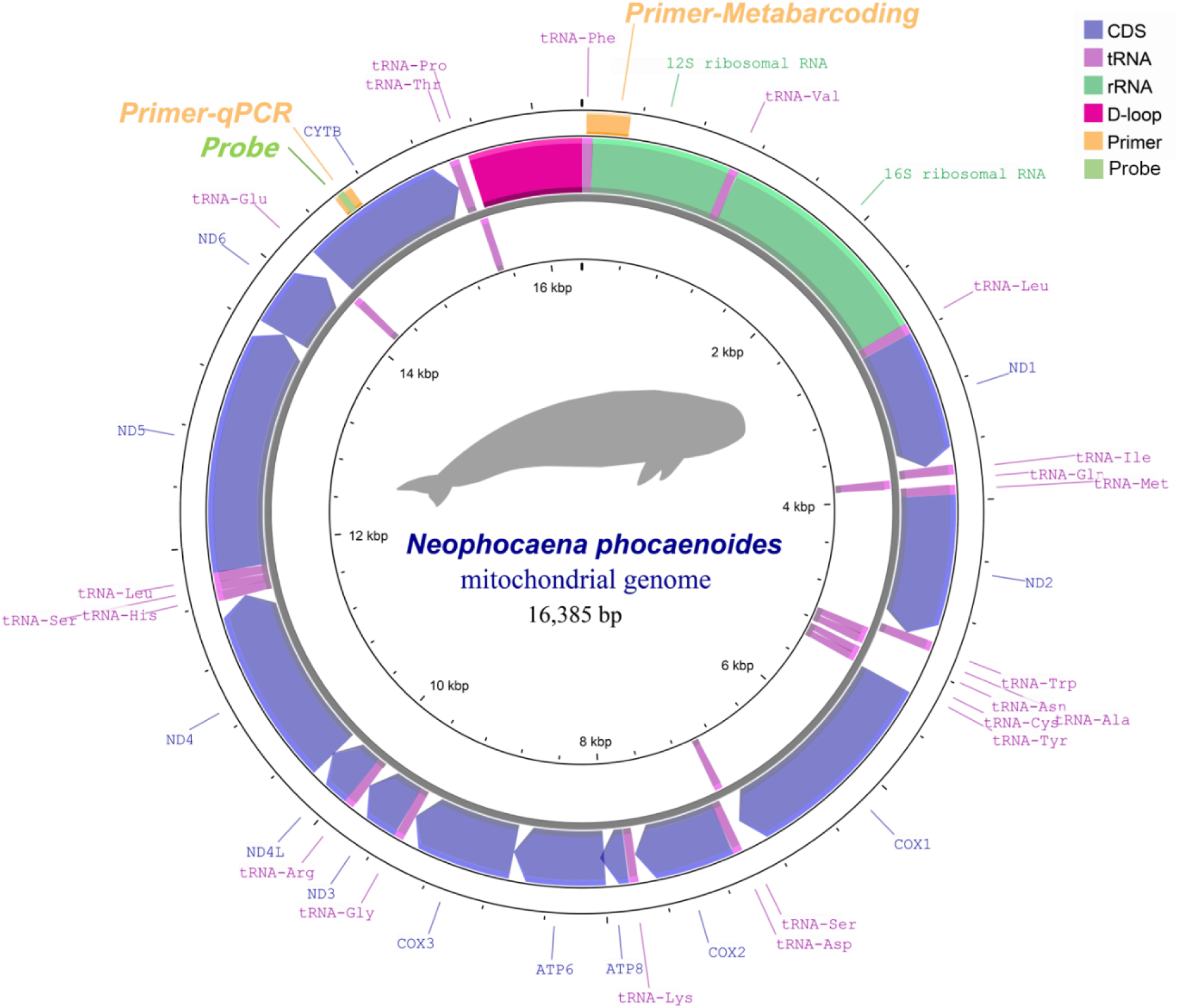
A Circos map of the mitochondrial genome of *Neophocaena phocaenoides* and the positions of the target regions of the qPCR and metabarcoding. Yellow color indicates the target regions of the qPCR and metabarcoding primers, while light-green color indicates the position of the qPCR probe.

For eDNA metabarcoding, we used a recently developed cetacean-specific primer pair, µCeta [36]. µCeta targets a 267-bp DNA fragment of the mitochondrial 12S region (Table 1 and Fig. 2). The performance of the µCeta primer pair was previously validated, and it detected resident cetaceans (*S. chinensis* and *N. phocaenoides*) in Hong Kong water.

### 2.3 Determination of the Limit of Detection (LOD) and Limit of Quantification (LOQ) of previously developed qPCR primers and probe

Before analyzing field samples, we evaluated the following four properties of the qPCR primers and probe: the intraspecific variation of the target region, the specificity, the limit of detection (LOD), and the limit of quantification (LOQ).

First, to examine if there are any intraspecific variations in the target region, we used tissue-extracted DNA. DNA of the Indo-Pacific finless porpoise was extracted from muscle tissue samples preserved at OPHK using the DNeasy Blood & Tissue Kit (Qiagen, Hilden, Germany) following the manufacturer’s protocol. Then, we amplified our target region (MT-CYTB) of the tissue-extracted DNA using the following PCR conditions: 20 µL, containing 10 µL of Platinum SuperFi II Master Mix (Thermo Fisher Scientific, Waltham, MA, USA), 2 µL of the forward and reverse primer (each 5 µM) (forward primer = ACGGCTCCTACATATTCCAAGAAAC; reverse primer = TGGGAGAATAAAGTGGAAGGCG; amplicon length = 248 bp), 4 µL of nuclease-free water and 2 µL of tissue-extracted DNA. We sent the PCR product to a company for Sanger sequencing.

Regarding the specificity of primers, we utilized the Primer-BLAST online tool (https://www.ncbi.nlm.nih.gov/tools/primer-blast/) to check if this primer could amplify the target region of other species. The result showed that this primer did not match any closely related species. Furthermore, we checked the amplification using synthesized DNA of *S. chinensis*, as it is also a resident species in Hong Kong waters. The amplification was tested on Thermo Fisher QuantStudio^TM^ 6 Flex using a 20 µL reaction volume containing 10 µL of TaqMan Environmental Master Mix 2.0 (Thermo Fisher Scientific), 2 µL of primer-probe mix (each 9 µM F/R qPCR primer and 1.25 µM probe), 0.2 µl of AmpErase Uracil N-glycosylase (UNG) (Thermo Fisher Scientific), 5.8 µL of nuclease-free water, and 2 µL of template DNA. The thermal cycle profile was as follows: 50°C for 2 min (UNG activation), 95°C for 10 min (denaturation), followed by 55 cycles of 95°C for 15 sec and 60°C for 1 min (annealing/extension). Fluorescence was measured at the 60°C step. We confirmed that *S. chinensis* DNA was not amplified by the primers.

Regarding the LOD and LOQ, we conducted qPCR using synthetic DNA of the Indo-Pacific finless porpoises to determine them. There are several possible definitions of the LOD and LOQ, and in this study we follow the definitions of Klymus et al. [39]. Specifically, the LOD was the lowest DNA concentration yielding ≥ 95% positive detection among technical replicates, and the LOQ was the lowest DNA concentration quantifiable with a coefficient of variation (C.V.) ≤ 35%. For robust LOD determination, at least 20 replicates per each DNA concentration are required. We obtained synthesized DNA (128 bp fragment in the MT-CYTB region) as standard DNA and prepared 24 replicates for each DNA concentration (DNA concentration ranged from 0 to 10^5^ copies/µL). For the 24 replicates of each DNA concentration, qPCR was performed using the conditions described above.

To calculate the LOD and LOQ, we utilized a custom R script provided by Klymus et al. [39], which determined a “modeled LOD” by selecting the best-fit model (from asymptotic regression, log-logistic, Michaelis-Menten, and Weibull I/II) for the concentration-detection probability relationship. The “modeled LOQ” was also determined by modeling the concentration-C.V. relationship. Details are provided by Klymus et al. [39] and in the associated GitHub repository (https://github.com/cmerkes/qPCR_LOD_Calc).

### 2.4 Study area, water sampling, and the measurement of environmental variables

According to a previous study of the distribution of the Indo-Pacific finless porpoise in Hong Kong waters [36–37], one of its major habitats is in the eastern and southern areas of Hong Kong waters, and thus, water samples were collected from areas surrounding the Soko Islands in the southeastern Hong Kong waters (Fig. 3). The survey was conducted on 25^th^ March 2025. We collected 1 L of surface (about 20 cm deep) and bottom (1 m above the bottom) water samples from 20 locations, which resulted in a total of 40 water samples (Fig. 3). The water depth in this region is generally shallow (less than about 20 m) [40]. In addition to the real field water samples, we filtered 1 L of MilliQ water samples as “field negative controls” at the three sampling locations (the first, middle, and final sampling locations). All of the samples, including negative control samples, were filtered using Sterivex filter cartridges (φ0.45-µm, SVHV010RS; Merck Millipore, Darmstadt, Germany) on the boat using the gravity filtration method [41]. Then, 2 ml of RNAlater (Thermo Fisher Scientific) was added to the filter cartridges, which were kept in a cooler box until they were brought back to the lab. Then, the samples were stored at –20°C in the lab until DNA extraction. Water temperature, salinity, and dissolved oxygen (DO) of sampling locations were measured using a CTD sensor (Model ASTD102; JFE Advantech Co., Ltd., Nishinomiya, Hyogo, Japan).

**Fig. 3.**
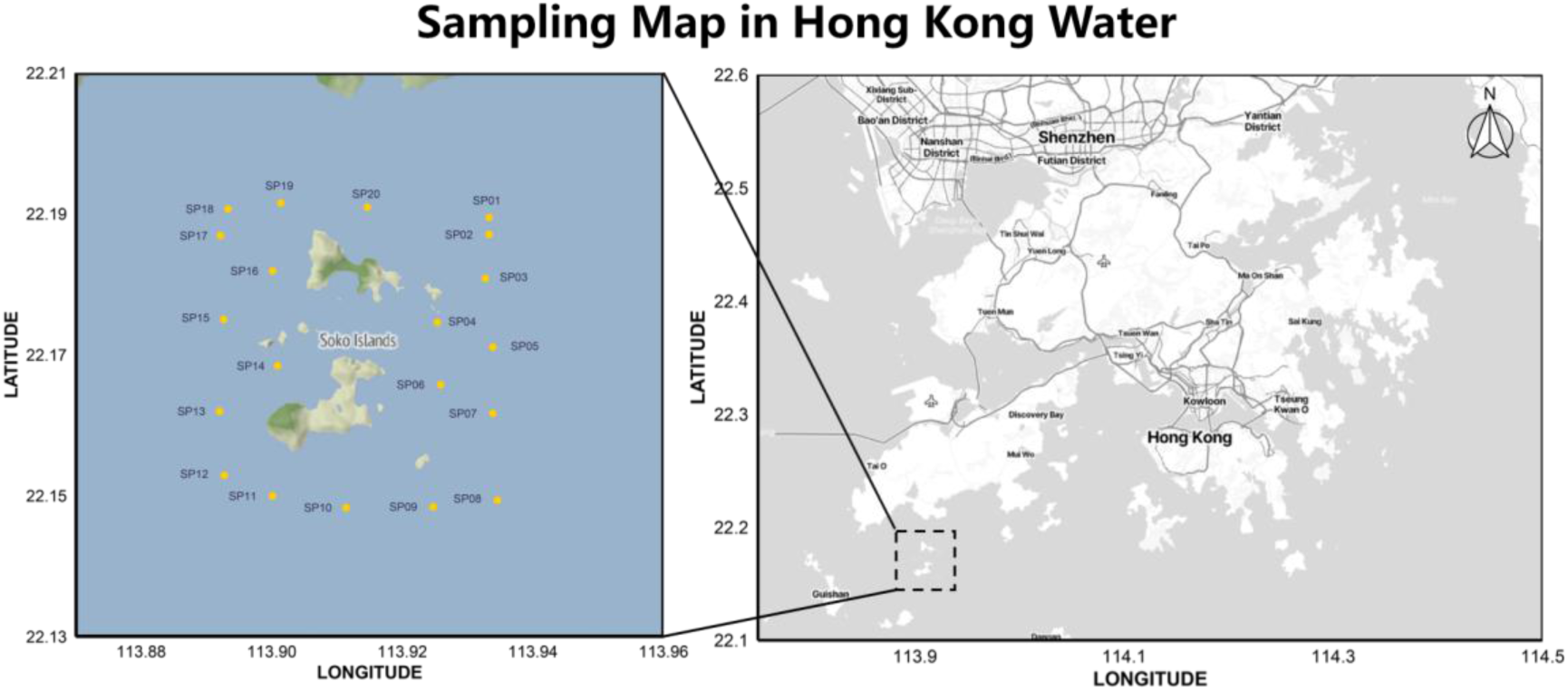
Sampling map of field survey in Hong Kong water. (Right) Hong Kong waters. The rectangle with dashed borders indicates our survey area. (Left) An enlarged map of our survey area. Each point indicates a sampling site for both surface and bottom water samples. Maps created using the ggmap package of R; ©Stadia Maps, ©Stamen Design, ©OpenMapTiles and ©OpenStreetMap.

### 2.5 eDNA extraction from Sterivex filter cartridges

eDNA was extracted from the Sterivex filter cartridges, including 40 field water samples and three negative control samples, using a protocol described previously [42] using the DNeasy Blood & Tissue kit. Briefly, RNAlater was removed from the filter cartridge by vacuum filtration, followed by a wash using 1 ml of MilliQ water. Then Lysis Mix, consisting of Buffer ATL (380 μL) and Proteinase K (20 μL), was added into each filter cartridge. The materials on the cartridge filters were subjected to cell lysis by incubating the filters at 56°C for 30 min. After the incubation, Buffer AL (400 μL) and 99.5% EtOH (400 μL) were added into each filter cartridge, followed by an additional 10-min incubation at 56°C. Lysate was transferred to a new 2-ml tube from the filter cartridge using a manual centrifuge (Handzentrifuge, Hittich, Westphalia, Germany). The collected DNA was purified using a DNeasy Blood & Tissue kit following the manufacturer’s protocol. DNA was eluted using 100 µL Buffer AE, and extracted DNA was stored at –20°C.

### 2.6 Detection of Indo-Pacific finless porpoise eDNA using qPCR

The qPCR was performed as described in section 2.2: Each 20-µL PCR reaction mix was prepared using Environmental Master Mix 2.0 (10 µL), AmpErase UNG (0.2 µL), primer-probe mix (2 µL), nuclease-free water (5.8 µL), and 2.0 µL eDNA template. Synthesized standard DNAs (0–10^5^ copies/µL) and samples were run in triplicate on a 96-well plate. Thermal cycle profile was the same as described above. The DNA copy numbers in water samples were calculated using the standard curve created with their C_t_ values. Using values from the three replicates for each sample, we calculated the average concentration as the representative value for the sample. If the calculated concentration of each replicate was below LOD, we regarded it as “not detected” and removed it from the calculation.

### 2.7 Detection of Indo-Pacific finless porpoise using eDNA metabarcoding

For eDNA metabarcoding, µCeta primer set was utilized [36]. Briefly, DNA libraries were prepared using a two-step “early-pooling” PCR protocol [43] (Fig. S1). The first-round PCR was performed in 20-μL reactions containing 10 μL of 2 × Platinum SuperFi II PCR Master Mix, 2 μL each of 5 μM µCeta-F/R primers with tag sequences (Table 1), 2 μL of nuclease-free water, and 4 μL of template DNA. The thermal cycle profile consisted of initial denaturation at 98°C for 30 s, followed by 35 cycles of 98°C for 10 s, 60°C for 10 s, and 72°C for 15 s, with a final extension at 72°C for 5 min. PCR negative controls (nuclease-free water was used instead of eDNA sample) were included to monitor contamination throughout the workflow. PCR products were purified using AMPure XP beads (Beckman Coulter, Brea, CA, USA) (PCR product: AMPure XP = 1:0.8).

The second-round PCR added dual indices in 20-μL reactions containing 10 μL of 2 × Platinum SuperFi II PCR Master Mix, 2 μL each of 5 μM Illumina 2^nd^ PCR primers (Table 1), 4 μL of nuclease-free water and 2 μL of purified first PCR product, using the following conditions: initial denaturation at 98°C for 30 s, 10 cycles of 98°C for 10 s and 72°C for 15 s, and final extension at 72°C for 5 min. Following the second-round PCR, the indexed PCR products were pooled, and the pooled product was purified using AMPure XP. The purified DNAs with the target size were excised using the E-Gel SizeSelect (Thermo Fisher Scientific). The library was sequenced on the Illumina iSeq100 system (Illumina, San Diego, CA, USA) using iSeq100 Reagent v2 (1 × 300 bp SE) with 30% PhiX spike-in. Two sequencing runs were performed to analyze all of our water samples.

### 2.8 Sequence data analysis of eDNA metabarcoding data

Sequencing data was demultiplexed using Illumina BaseSpace and a custom script with Cutadapt v4.7 [44]. Raw sequences were quality-filtered using fastp v0.23.4 [45] to remove low-quality reads and adapter sequences, followed by primer trimming with Cutadapt. The processed reads were then analyzed using DADA2 [46] implemented in R [47] to generate amplicon sequence variants (ASVs). DADA2 processing involved quality filtering, error rate estimation, dereplication, error correction, read merging, and chimera removal. These processes were performed separately for each iSeq run. We merged ASV tables created by the two iSeq runs using the *mergeSequenceTables* function. After merging the results from multiple runs, ASVs were further clustered into operational taxonomic units (OTUs) at 97% similarity using the DECIPHER package in R [48].

Taxonomic assignment process was performed using “Claident” v0.9.2021.10.22 (https://www.claident.org/), which implements a conservative approach [49]. For OTUs generated through 97% similarity clustering, we conducted taxa assignments using two databases: the *overall_genus* reference database and the *overall_class* database. The *overall_genus* database contains NCBI nt sequences which have genus or lower-level taxonomic information, while the *overall_genus* database contains NCBI nt sequences which have class or lower-level taxonomic information. Each analysis involved three sequential steps: database caching (*clmakecachedb*), sequence identification (*clidentseq*), and taxonomic assignment (*classigntax*). The results were then merged using *clmergeassign* with descending priority. Also, we used a manual BLAST search (https://blast.ncbi.nlm.nih.gov/Blast.cgi) to check the taxa assignment results.

For subsequence analyses, samples with fewer than 10 sequence reads were excluded (i.e., most negative controls and some field samples). In this step, we skipped the “rarefaction” process, as the number of detected OTUs was small (a few cetacean OTUs) partly because of the high specificity of µCeta primers.

### 2.9 Statistical analysis

All statistical analyses were performed using R v4.4.3 [47]. The effects of environmental parameters on the eDNA detection were tested using the generalized linear models (GLMs), and the effects of layers (i.e., surface or bottom) were tested using the chi-square test. In addition, we tested the effects of the two eDNA approaches (qPCR and metabarcoding) on the eDNA detection using the test for equality of proportions. The relative abundance data and correlation data were visualized using the *ggplot2* package in R [50].

## 3. Results

### 3.1 Intraspecific Variations of the target region, and the Specificity, LOD, and LOQ of qPCR primers and probe

Sangar sequencing of the qPCR target region of the tissue-extracted DNA showed no intraspecies difference among finless porpoise individuals in Hong Kong waters. In addition, our qPCR assay confirmed the amplification of tissue-extracted DNA of the finless porpoises. On the other hand, DNA extracted from the Indo-Pacific humpback dolphin was not amplified by the qPCR.

The LOD and LOQ were determined by the analysis of synthesized standard DNA. For LOD, the best fit curve was Weibull type II, and the estimated modeled LOD was 3 copies per reaction (equivalent to 1.5 copies/µL in template DNA, as we used 2 µL of template DNA). For LOQ, the best fit curve was the 5th-order polynomial model, and the estimated modeled LOQ was 34 copies per reaction (equivalent to 17 copies/µL in template DNA) (Fig. S2).

### 3.2 Detection of Indo-Pacific finless porpoise eDNA using qPCR

The species-specific qPCR assay successfully detected finless porpoise eDNA at multiple locations around the Soko Islands (Fig. 4A). Specifically, finless porpoise eDNA was detected (i.e., eDNA concentrations above LOD) in 5 out of 20 surface water samples and 3 out of 20 bottom water samples (Table S1). Unfortunately, all of the detected eDNA concentrations fell below LOQ (34 copies per reaction, equivalent to 1.7 × 10^3^ copies/L seawater). Thus, only the qualitative results (i.e., detected or not) are reported. Among locations with positive detection, SP01 exhibited the most consistent detections, yielding positive detections (eDNA concentrations above LOD) in five of six technical replicates of qPCR from the surface and bottom layers. Among other locations, only SP11 also yielded positive detections in the surface and bottom layers, but the detections were less consistent than thoseat SP01 (i.e., detection depends on technical replicates and water layers). Overall, the proportions of positive eDNA detection were 25% for the surface water samples and 15% for the bottom water samples (Fig. 4A).

**Fig. 4.**
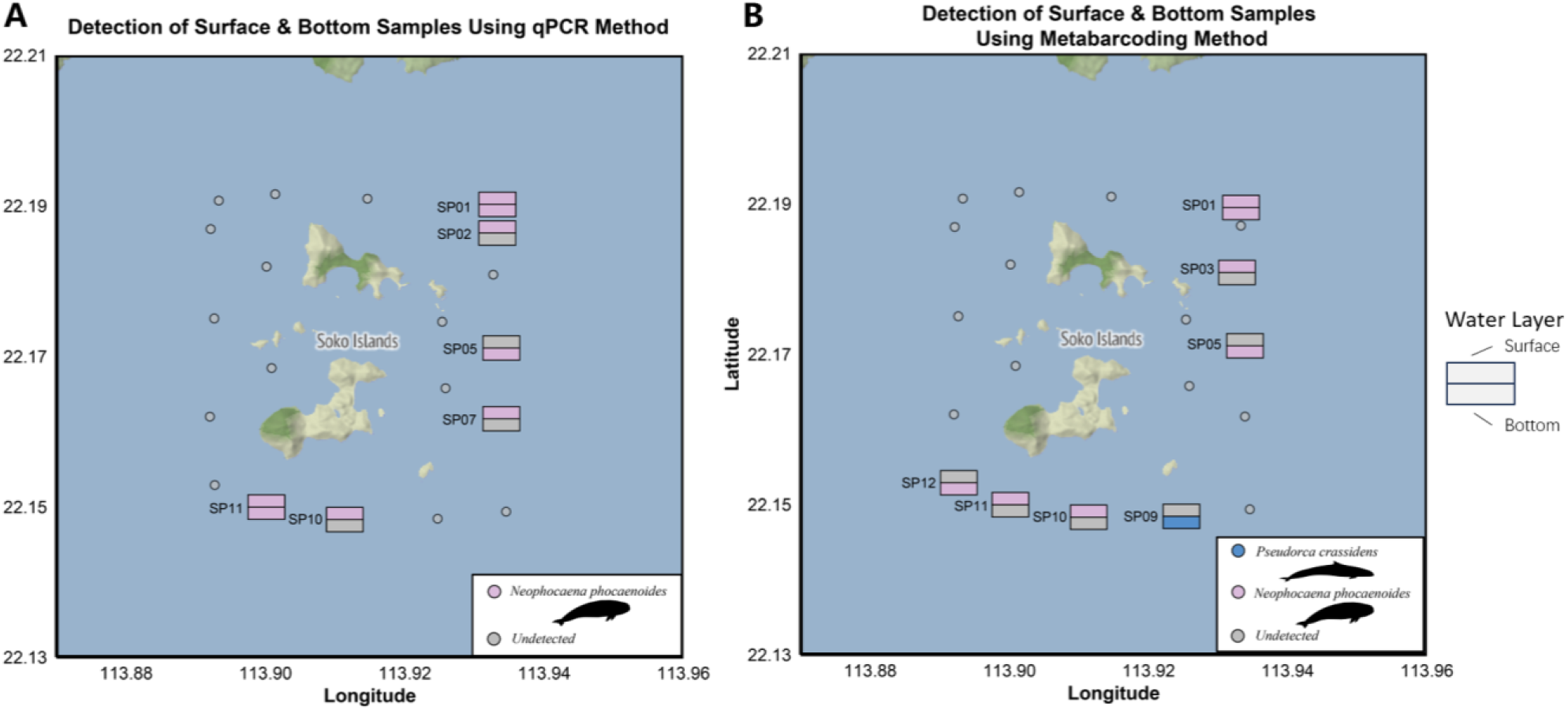
Detection of eDNA of Indo-Pacific finless porpoises and other cetaceans. (**A**) qPCR-based detection of Indo-Pacific finless porpoises’ eDNA for surface and bottom water samples. The upper block indicates surface sample; the lower one indicates bottom sample. Pink, color-filled blocks indicate positive detections, while gray-filled ones indicate no detection. Note that only the qualitative results are shown here, as all the values were below the limit of quantification (LOQ). (**B**) Metabarcoding-based detection of cetacean eDNA for surface and bottom water samples. The upper block indicates surface sample; the lower one indicates bottom sample. The presence of a species is indicated by a filled block, with colors representing different cetacean species (see the legends inside the figures). A grey block indicates that no cetacean eDNA was detected. Cetacean silhouette credit: finless porpoise (by Chris huh; CC BY-SA 3.0); *Pseudocrca crassidens*, false killer whale (by Chris huh; CC BY-SA 3.0).

### 3.3 Detection of Indo-Pacific finless porpoise eDNA using eDNA metabarcoding

Our µCeta eDNA metabarcoding generated a total of 606,588 sequence reads from 49 samples (including negative control samples), with an overall >Q30 score of 90.2%. After quality filtering and DADA2 denoising, 580,976 reads remained (95.77% of the total reads).

After the OTUs clustering process, 17 OTUs were detected. Among them, we detected two cetacean OTUs, which contained a total of 565,579 sequence reads (97.35 % of DADA2 processed reads) in all water samples. One OTU, which was assigned to the genus *Neophocaena* (finless porpoise), was detected from multiple locations and layers. Given that the Indo-Pacific finless porpoise is the only species of this genus known to inhabit Hong Kong waters, we attribute this OTU to the Indo-Pacific finless porpoise.

The µCeta metabarcoding successfully detected the Indo-Pacific finless porpoise eDNA in four surface and three bottom water samples (Fig. 4B and Table S2). The location SP01 was the only location where the porpoise eDNA was detected in both the surface and bottom layers by µCeta metabarcoding. The other OTU, which was assigned to the false killer whale (*Pseudorca crassidens*), was detected from a single location, SP09 (Fig. 4B). No eDNA sequence reads of the Indo-Pacific humpback dolphin were detected from the water samples.

We detected some non-cetacean reads (in total, 8,901 reads, which corresponded to 1.57 % of cetacean eDNA; Table S3), and they could not be assigned to a certain genus or class. After performing BLAST-search manually, we concluded that some of them may belong to eukaryotic microorganisms, but their lower taxonomic identity could not be identified. From negative control samples, we detected almost non-cetacean reads only (Tables S2–S3). Also, the number of non-cetacean reads in the negative control samples was negligible (totally 1.12% of all the reads after quality filtering and DADA2 denoising, including 0.43% from three field negative controls, 0.21% from two PCR negative controls, and 0.47% from four DNA extraction negative controls; Table S3), showing that there was no severe contamination. These metrics confirm the high quality and reliability of the sequencing data.

### 3.4 The effects of environmental parameters, water layers, and eDNA analysis methods on the eDNA detection

Environmental parameters had little influence on eDNA detection (Fig. 5). In the qPCR analysis, in general, dissolved oxygen (DO) and salinity showed a negative relationship with finless porpoise eDNA detection, while temperature exhibited a positive relationship with the eDNA detection. However, the effects of environmental parameters on the eDNA detections were not statistically significant (GLM, *p* > 0.05). In the metabarcoding analysis, water temperature showed a positive relationship with the eDNA detection, while salinity and DO showed a negative relationship, but these relationships were also not statistically significant (GLM, *p* > 0.05).

**Fig. 5.**
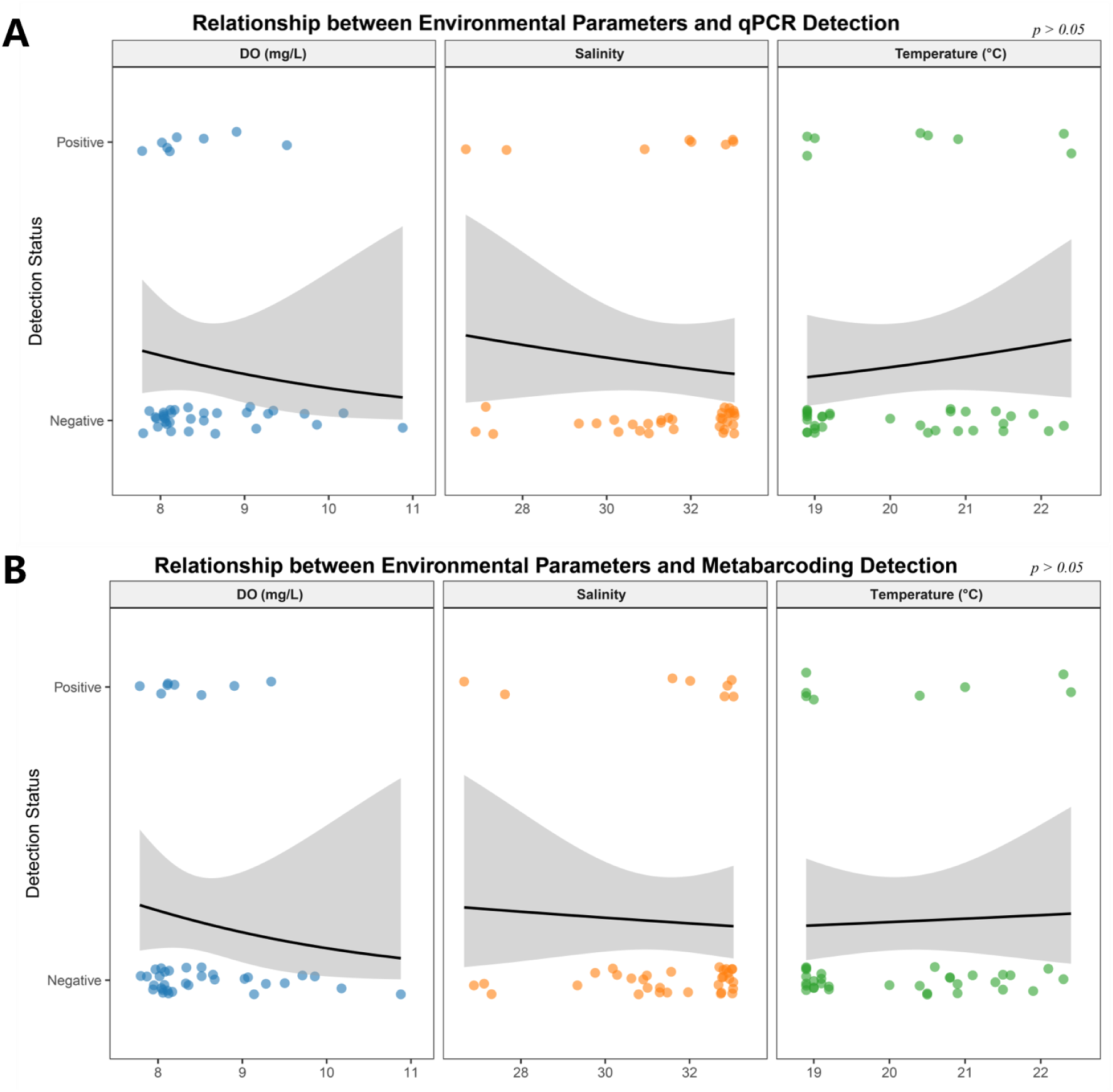
The relationships between environmental factors and Indo-Pacific finless porpoise eDNA detection. (**A**) The relationship between qPCR-based eDNA detection and three environmental factors: DO, salinity, and temperature (B) The relationship between metabarcoding-based eDNA detection and the three environmental factors: DO, salinity, and temperature. Note that points are slightly jittered along the *y*-axis for better visibility. Solid black lines represent the predicted relationship using a binomial generalized linear model (GLM), and grey shaded areas indicate 95% confidence intervals. All correlations were insignificant (*p* > 0.05).

We also compared the detection rates of the Indo-Pacific finless porpoise eDNA between different water layers (i.e., surface versus bottom layers) and methods (i.e., qPCR versus metabarcoding). In general, surface water samples exhibited slightly higher detection rate, and both methods detected eDNA more frequently from surface water samples than from bottom water samples (Fig. 6A, B). This tendency was slightly more pronounced in the qPCR results, but there were no statistically significant differences in the detection rate between the two layers for either method (equality of proportion test, *p* > 0.05; Fig. 6A, B).

**Fig. 6.**
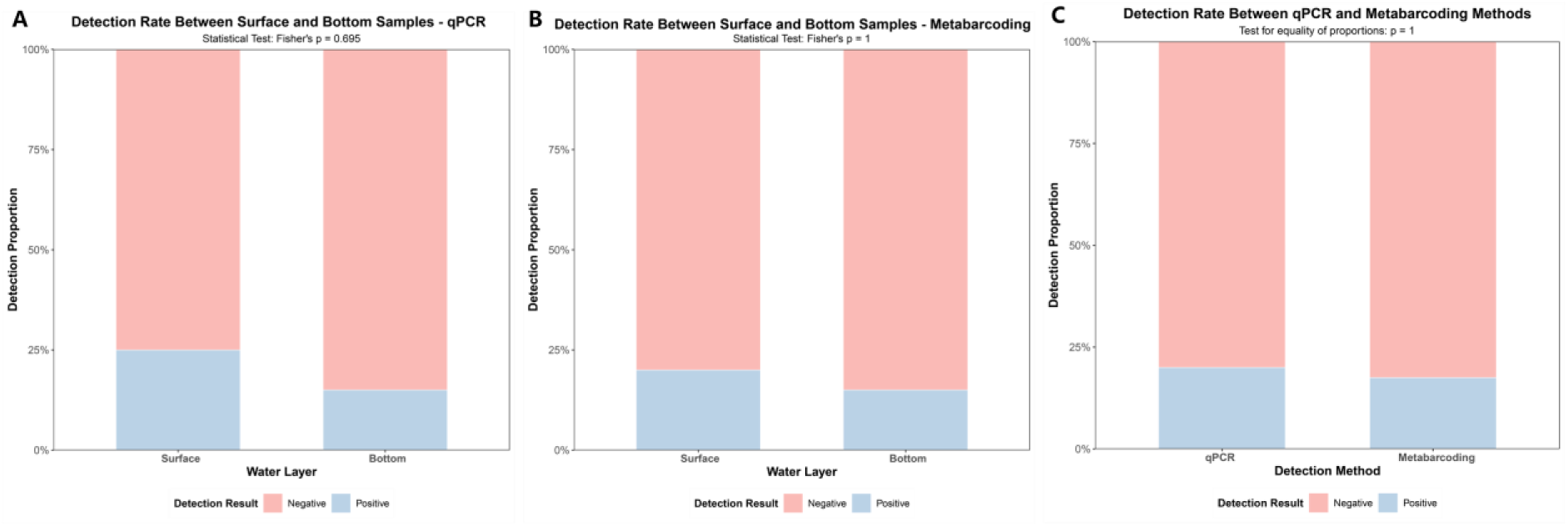
The relationship of the Indo-Pacific finless porpoise eDNA detection rates among different water layers and eDNA detection methods. (**A**) eDNA detection rates of surface and bottom water samples using qPCR. (B) eDNA detection rates of surface and bottom water samples using metabarcoding. (C) eDNA detection rate of the qPCR and metabarcoding methods.

Regarding the difference between the qPCR and metabarcoding, the detection rate was approximately the same (qPCR detection rate = 20%; metabarcoding detection rate = 17.5%). The detection rate was not statistically significantly different between the two methods (equality of proportion test, *p* > 0.05; Fig. 6C). In total, five samples were “positive” by both methods, while the other six samples were “positive” by one of the two methods (Fig. 4).

## 4. Discussion

### 4.1 Specificity, LOD and LOQ of the qPCR primers and probe

The qPCR target region we utilized in this study showed no intraspecific variation among the Indo-Pacific finless porpoise individuals stranded in Hong Kong coastal areas, confirming its applicability across different individuals. The assay successfully amplified tissue-derived DNA of the finless porpoise but not that of the Indo-Pacific humpback dolphin, supporting the specificity of the qPCR primers and probe.

The modeled LOD and LOQ were 3 and 34 copies per reaction, respectively (Fig. S2). Such sensitivity is critical for reliable qualitative detection and quantification of target eDNA in environmental samples [39, 51–52]. When compared to previously reported qPCR assays of eDNA, the LOD and LOQ of our assay are comparable. For example, the multi-laboratory assessment by Klymus et al. (2020) evaluated 36 different qPCR assays of eDNA, and the modeled LOD values ranged from 2.19 to 260 copies per reaction, and the modeled LOQ values spanned from 6 to 839 copies per reaction [39]. The LOD of our qPCR assay was 3 copies/reaction, which was well within the lower range of the reported values. Also, the LOQ of our qPCR assay was 34 copies/reaction, which was lower than that of the majority of assays in Klymus et al. [39]. These results suggest that our qPCR assay is reasonably sensitive for facilitating future field-based eDNA monitoring of the Indo-Pacific finless porpoise.

### 4.2 Detection of the finless porpoise eDNA using qPCR

The qPCR assay detected finless porpoise eDNA in 8 out of 40 water samples (20%), but all positive samples showed eDNA concentrations below the LOQ, preventing reliable quantification. This detection rate is comparable to that reported in other cetacean eDNA studies using qPCR. For example, Hashimoto et al. (2024), utilizing the same primer set for finless porpoise detection in Osaka Bay, Japan, reported eDNA detection at 9 out of 50 sites (18%) [35]. The slightly higher detection rate in our study could reflect actual differences in porpoise abundance between Osaka Bay and Hong Kong waters, though variations in sampling strategy, water volume, and local hydrodynamic conditions may also contribute to these differences. Similarly, Kisero et al. (2025) reported detection rates of 21.4% (6/28) in summer and 53.6% (15/28) in winter for the Indo-Pacific humpback dolphins in Hong Kong waters using a qPCR approach [37–38]. It is important to note that detection rates are inherently influenced by the actual number and distribution of target species individuals in the survey area, making direct cross-study comparisons infeasible. The consistently low eDNA concentrations (e.g., below LOQ) were observed across all of the three studies, suggesting that cetacean eDNA in marine environments generally presents at very low concentrations, likely due to the cetaceans’ relatively low population densities and large home ranges.

In terms of horizontal distribution, the majority of positive eDNA detections tended to be localized from the northeast to east to southwest of the Soko Islands (Fig. 3). This weak spatial clustering could suggest potential aggregation of finless porpoise individuals in these areas during the sampling period. However, because this study represents a single sampling event, and because the spatial pattern was not statistically tested, the observed distribution pattern should be interpreted with caution.

### 4.3 Detection of eDNA using metabarcoding

Metabarcoding with the µCeta primer set detected two cetacean species: the Indo-Pacific finless porpoise and the false killer whale (Fig. 3). No non-cetacean vertebrate sequences were identified, confirming the high specificity of the µCeta primers. This may be advantageous for cetacean eDNA metabarcoding over earlier universal primers (e.g., Riaz12S [53], MiFish [54], and MarVer [55]) because co-amplification of fish and other taxa can reduce detection efficiency of low-abundance cetacean eDNA [55–57]. Among a total of 17 detected OTUs, two were assigned to cetaceans. The remaining 15 OTUs could not be definitively classified, though BLAST results suggested that some may represent eukaryotic microorganisms. Although we detected some non-target species, their sequence reads constituted a small proportion (1.57%) of cetacean eDNA reads, demonstrating the high specificity and efficiency of µCeta metabarcoding.

Similar to the qPCR results, finless porpoise eDNA was detected in the northeastern, eastern, and southern sampling locations (Fig. 3). Interestingly, false killer whale eDNA was detected in a bottom water sample collected at SP09. According to the AFCD record [37–38], false killer whales are one of the transient cetacean species in Hong Kong waters, and their eDNA detection could suggest that the false killer whale individuals were indeed there even though we did not directly sight them during the field survey. Further field surveys in different seasons and with broader spatial coverage will be required to investigate the spatial distribution patterns and the species occurrence of the transient species in Hong Kong waters.

### 4.4 The effects of environmental variables, water layers, and methods

No statistically significant correlations were observed between eDNA detection rates and dissolved oxygen (DO), salinity, or temperature with either the qPCR or metabarcoding approach (Fig. 5), suggesting that these environmental factors did not strongly influence eDNA detection within the spatiotemporal scale of our survey. This finding aligns with our previous research on the Indo-Pacific humpback dolphin eDNA in Hong Kong waters [37–38].

Both qPCR and metabarcoding showed comparable detection rates between surface and bottom water layers, with no statistically significant differences (Fig. 6). This may have been due to the shallow coastal setting and active physical forces such as wind and tides [58–59]. Under such conditions, eDNA released by cetaceans may be rapidly dispersed throughout the water column, reducing vertical stratification. These results may suggest that in similarly dynamic coastal environments, sampling at a single depth (e.g., surface water only) may suffice for detecting cetacean presence, thereby simplifying field protocols.

The species-specific qPCR assay and metabarcoding showed comparable detection rates for the finless porpoise, at 20% and 17.5%, respectively, with no statistically significant difference. The similar performance indicates that, in this study, both methods were equally effective in detecting the target species under field conditions. While the shorter target region of qPCR (128 bp vs. 267 bp in µCeta metabarcoding) may theoretically enhance the detection sensitivity [60], this was not pronounced in this initial survey. Interestingly, metabarcoding provided valuable supplementary information by detecting the false killer whale (*Pseudorca crassidens*) eDNA, demonstrating its utility for broader cetacean biodiversity screening [36].

### 4.5 Limitations and future perspectives

Although we successfully demonstrated that our qPCR and metabarcoding approaches effectively detected the resident finless porpoises and one transient species, the false killer whale, in Hong Kong waters, some limitations still remained. First, hydrological processes, such as currents, tides, and vertical mixing, which are known to govern the transport and persistence of eDNA in marine environments [61–62], were not explicitly integrated into our sampling design or analysis. These factors could significantly influence the spatial and temporal distribution of eDNA signals, thereby affecting the interpretation of detection patterns. Future studies can incorporate hydrodynamic modeling to account for the advection and dispersion of eDNA, which would enhance the ecological inferences derived from eDNA data and improve the accuracy of source estimation.

Second, the absence of independent monitoring data, such as systematic visual surveys or acoustic detections, limited our ability to validate eDNA-based occurrence records against actual animal presence or to estimate detection probability [63]. Future research would benefit from an integrated survey design that combines eDNA sampling with visual, acoustic, and telemetry methods. Such a multimodal framework would allow for a more robust evaluation of eDNA performance in detecting cetaceans and help to elucidate the relationships between eDNA signals and animal abundance or behavior.

Third, the low eDNA concentrations typically encountered in marine systems, as also reflected in our study, highlight the need for continued methodological advancements. A previous study showed that the concentration of marine eDNA after water filtration and extraction is often within the range of 0.01–1 copies/µL [64]. For such low-concentration eDNA, digital droplet PCR (ddPCR) may provide more robust detection and quantification [64]. Also, to improve the detection efficiency, the filtration of larger water volumes and optimization of DNA extraction protocols for low-biomass samples could be considered [60]. These refinements are essential to improve both the detection probability and the quantitative potential of eDNA-based assays for marine mammals.

Lastly, we propose several strategic directions to advance the field of eDNA-based cetacean monitoring [55]. Expanding eDNA surveys across broader spatial and temporal scales will be critical for establishing baseline distributions and understanding seasonal or interannual variability in cetacean occurrence. Furthermore, the integration of eDNA data with Species Distribution Models (SDMs) offers a powerful approach to elucidate habitat preferences and forecast marine species responses to environmental change [65–68]. By combining eDNA-derived occurrence data with spatially explicit environmental variables, such as sea surface temperature, bathymetry, and oceanographic fronts, future studies could predict suitable habitats and identify potential foraging or breeding grounds for cetacean conservation prioritization [69]. In addition, linking eDNA results from cetaceans with metabarcoding data of potential prey species could provide novel insights into trophic interactions and habitat functionality [34].

## 5. Conclusion

In this study, two eDNA approaches, species-specific qPCR and µCeta metabarcoding, successfully detected Indo-Pacific finless porpoises, demonstrating the potential of these approaches as reliable and non-invasive means for monitoring cetaceans in Hong Kong waters. The eDNA detections were relatively localized to the northeastern to eastern to southern areas of Soko Islands coastal regions. The absence of a statistically significant difference in the detection rates between surface and bottom water samples may have been due to vertically well-mixed finless porpoise eDNA, suggesting that more efficient water sampling could be done by focusing on the surface water samples only. qPCR and µCeta metabarcoding methods proved equally effective in detecting the resident finless porpoise in this study. Meanwhile, metabarcoding provided additional data through the detection of the false killer whale. Our study showed a basis for an eDNA-based cetacean monitoring framework in Hong Kong waters. By adopting an integrated and multidisciplinary framework, eDNA technology can substantially advance the conservation and management of elusive marine mammals, particularly in rapidly changing coastal ecosystems.

## Data and Code Accessibility

Illumina sequence data was deposited in NCBI (Accession BioProject No. = PRJNA1371113, BioSample No. = SAMN53489895 - SAMN53489930). Sanger sequence data was deposited in NCBI (GenBank Accession No. = BankIt3033257, PX703111 - PX703127)

## Acknowledgments

We thank Takamitsu OHIGASHI, Suixuan HUANG, Robinson O. Kisero, Chengbin, LIU, Lucia HU, Mingwei LI, Shuting Qiu, Ocean Rauszen ABBAS for their assistance in the field work and Leung Kwan Chak for his assistance in the tissue sample collection and management. We thank the Agriculture, Fisheries and Conservation Department (AFCD) of Hong Kong, and the Ocean Park Hong Kong (OPHK) and Foundation Hong Kong (OPCFHK) for their permission for using cetacean tissue samples and assistance in collecting them. This research was supported by The Hong Kong University of Science and Technology Startup Fund to MU, and GRF16100724 from the Research Grants Council of the Hong Kong SAR, China to MU.

## Author contributions

MU conceived research; XL and MU designed research; XL and MU validated qPCR primers and probe and performed field survey: XL performed experiments and data analysis with technical advice from MU; XL and MU wrote the manuscript.

## References

[1] Society for Marine Mammalogy. “Home.” Accessed November 20, 2025. http://marinemammalscience.org.

[2] L. Gilbert, T. Jeanniard-du-Dot, M. Authier, T. Chouvelon, J. Spitz, Composition of cetacean communities worldwide shapes their contribution to ocean nutrient cycling, NAT COMMUN, 14 (2023) 5823.

[3] Y.J. Sun, Shantou University, Shantou, 2022.

[4] J. De Weerdt, E.A. Ramos, E. Pouplard, M. Kochzius, P. Clapham, Cetacean strandings along the Pacific and Caribbean coasts of Nicaragua from 2014 to 2021, Marine biodiversity records, 14 (2021) 1.

[5] S.M. Maxwell, E.L. Hazen, S.J. Bograd, B.S. Halpern, G.A. Breed, B. Nickel, N.M. Teutschel, L.B. Crowder, S. Benson, P.H. Dutton, H. Bailey, M.A. Kappes, C.E. Kuhn, M.J. Weise, B. Mate, S.A. Shaffer, J.L. Hassrick, R.W. Henry, L. Irvine, B.I. McDonald, P.W. Robinson, B.A. Block, D.P. Costa, Cumulative human impacts on marine predators, NAT COMMUN, 4 (2013) 2688–2688.

[6] T.A. Jefferson, S.K. Hung, Neophocaena phocaenoides, Mammalian Species, 746 (2004) 1–12.

[7] T.A. Jefferson, J.E. Moore, Abundance and Trends of Indo-Pacific Finless Porpoises (*Neophocaena phocaenoides*) in Hong Kong Waters, 1996–2019, FRONT MAR SCI, 7 (2020).

[8] L. Fang, M. Li, X. Wang, Y. Chen, T. Chen, Indo-Pacific finless porpoises presence in response to pile driving on the Jinwan Offshore Wind Farm, China, FRONT MAR SCI, 10 (2023).

[9] M. Nithyanandan, Y. Bohadi, Incidental mortality events of the Indo-Pacific Finless Porpoise, *Neophocaena phocaenoides* in Kuwait, Northwestern Arabian Gulf, ZOOL MIDDLE EAST, 67 (2021) 373–376.

[10] T.A. Jefferson, S.K. Hung, L. Law, M. Torey, Distribution and abundance of finless porpoises in Hong Kong and adjacent waters of China, The Raffles Bulletin of Zoology, (2002).

[11] T.A. Jefferson, S.K. Hung, An updated, annotated checklist of the marine mammals of Hong Kong, MAMMALIA, 71 (2007).

[12] J. Watts, Further observations on the hydrology of the Hong Kong territorial waters, Hong Kong Fisheries Bulletin, 3 (1973) 9–35.

[13] B. Morton, G. Blackmore, South China Sea, MAR POLLUT BULL, 42 (2001) 1236–1263.

[14] J.H.W. Lee, P.J. Harrison, C. Kuang, K. Yin, Eutrophication Dynamics in Hong Kong Coastal Waters: Physical and Biological Interactions, The Environment in Asia Pacific Harbours, Springer Netherlands, Dordrecht, 2006. pp. 187–206.

[15] W. Ng, F.C.C. Leung, S.T.C. Chak, G. Slingsby, G.A. Williams, Temporal genetic variation in populations of the limpet Cellana grata from Hong Kong shores, MAR BIOL, 157 (2010) 325–337.

[16] K.W. Li, H.Y. Mok, LONG TERM TRENDS OF THE REGIONAL SEA LEVEL CHANGES IN HONG KONG AND THE ADJACENT WATERS, WORLD SCIENTIFIC2011. pp. 349–359.

[17] T.A. Jefferson, Population Biology of the Indo-Pacific Hump-Backed Dolphin in Hong Kong Waters Author(s): Thomas A. Jefferson Reviewed work(s): Source: Wildlife Monographs, No. 144, Population Biology of the Indo-Pacific Hump-Backed Dolphin in Hong Kong Waters (Oct., 2000), pp. 1–65 Published by: Allen Press Stable URL: http://www.jstor.org/stable/3830809. Accessed: 16/12/2012 14:42, (2000).

[18] IUCN, Indo-Pacific Finless Porpoise, 2025. https://www.iucnredlist.org/species/198920/50386795

[19] X. Zhang, W. Lin, R. Yu, X. Sun, Y. Ding, H. Chen, X. Chen, Y. Wu, Tissue partition and risk assessments of trace elements in Indo-Pacific Finless Porpoises (*Neophocaena phocaenoides*) from the Pearl River Estuary coast, China, CHEMOSPHERE, 185 (2017) 1197–1207.

[20] W. Lin, L. Karczmarski, J. Li, S.C.Y. Chan, L. Guo, Y. Wu, Differential population dynamics of a coastal porpoise correspond to the fishing effort in a large estuarine system, AQUAT CONSERV, 29 (2019) 223–234.

[21] C. Boyd, R.C. Hobbs, A.E. Punt, K.E.W. Shelden, C.L. Sims, P.R. Wade, Bayesian estimation of group sizes for a coastal cetacean using aerial survey data, MAR MAMMAL SCI, 35 (2019) 1322–1346.

[22] P.S. Hammond, C. Lacey, A. Gilles, S. Viquerat, P. Boerjesson, H. Herr, K. Macleod, V. Ridoux, M. Santos, M. Scheidat, J. Teilmann, J. Vingada, N. Oeien, Estimates of cetacean abundance in European Atlantic waters in summer 2016 from the SCANS-III aerial and shipboard surveys, Wageningen Marine Research2017.

[23] D. Gillespie, D.K. Mellinger, J. Gordon, D. McLaren, P. Redmond, R. McHugh, P. Trinder, X.Y. Deng, A. Thode, PAMGUARD: Semiautomated, open source software for real-time acoustic detection and localization of cetaceans., J ACOUST SOC AM, 125 (2009) 2547–2547.

[24] V.M. Janik, Cetacean vocal learning and communication, CURR OPIN NEUROBIOL, 28 (2014) 60–65.

[25] P. Taberlet, E. Coissac, L.H.R. Mehrdad Hajibabaei, Environmental DNA, MOL ECOL, (2012).

[26] E.E. Sigsgaard, I.B. Nielsen, S.S. Bach, E.D. Lorenzen, D.P. Robinson, S.W. Knudsen, M.W. Pedersen, M.A. Jaidah, L. Orlando, E. Willerslev, P.R. Møller, P.F. Thomsen, Population characteristics of a large whale shark aggregation inferred from seawater environmental DNA, NAT ECOL EVOL, 1 (2016) 0004.

[27] S. Tsuji, N. Shibata, R. Inui, R. Nakao, Y. Akamatsu, K. Watanabe, Environmental DNA phylogeography: Successful reconstruction of phylogeographic patterns of multiple fish species from cups of water, MOL ECOL RESOUR, 23 (2023) 1050–1065.

[28] M.L. Rourke, A.M. Fowler, J.M. Hughes, M.K. Broadhurst, J.D. DiBattista, S. Fielder, J. Wilkes Walburn, E.M. Furlan, Environmental DNA (eDNA) as a tool for assessing fish biomass: A review of approaches and future considerations for resource surveys, Environmental DNA, 4 (2022) 9–33.

[29] T. Takahara, T. Minamoto, H. Yamanaka, H. Doi, Z. Kawabata, J.A. Gilbert, Estimation of fish biomass using environmental DNA, PLOS ONE, 7 (2012) e35868–e35868.

[30] S. Yang, Y. Jin, S. Li, Z. Liu, Integrated approaches for comprehensive cetacean research and conservation in the East China Sea, MAR POLLUT BULL, 206 (2024) 116789.

[31] L.R. Harper, L. Lawson Handley, A.I. Carpenter, M. Ghazali, C. Di Muri, C.J. Macgregor, T.W. Logan, A. Law, T. Breithaupt, D.S. Read, A.D. McDevitt, B. Hänfling, Environmental DNA (eDNA) metabarcoding of pond water as a tool to survey conservation and management priority mammals, BIOL CONSERV, 238 (2019) 108225.

[32] R.O. Kisero, S. Tsuji, T. Ohigashi, L. Porter, E. Matrai, M. Ushio, Detection of environmental DNA of the Indo-Pacific humpback dolphins in Hong Kong waters using quantitative PCR, bioRxiv, (2025) 2025.10.30.685690.

[33] G. Boldrocchi, L. Conte, P. Galli, R. Bettinetti, E. Valsecchi, Cuvier’s beaked whale (*Ziphius cavirostris*) detection through surface-sourced eDNA: A promising approach for monitoring deep-diving cetaceans, ECOL INDIC, 161 (2024) 111966.

[34] S. Zhang, Y. Cao, B. Chen, P. Jiang, L. Fang, H. Li, Z. Chen, S. Xu, M. Li, Assessing the potential use of environmental DNA for multifaceted genetic monitoring of cetaceans: Example of a wandering whale in a highly disturbed bay area, ECOL INDIC, 148 (2023) 110125.

[35] N. Hashimoto, T. Iwata, N. Kihara, K. Nakamura, M.K. Sakata, T. Minamoto, Detection of environmental DNA of finless porpoise (*Neophocaena asiaeorientalis*) in Osaka Bay, Japan, CONSERV GENET RESOUR, (2024).

[36] M. Ushio, S. Ozawa, S. Oka, T. Sado, R.O. Kisero, L. Porter, E. Matrai, M. Miya, μCeta: A Set of Cetacean-Specific Primers for Environmental DNA Metabarcoding With Minimal Amplification of Non-Target Vertebrates, Environmental DNA, 7 (2025) e70193.

[37] H.K.C.R. Project, Monitoring of Marine Mammals in Hong Kong Waters (2022-23): Final Report. Hong Kong Cetacean Research Project Report to the Agriculture, Fisheries and Conservation Department, Hong Kong Cetacean Research, Hong Kong, 2023.

[38] S.K. Hung., J.Y. Wang, Monitoring of Marine Mammals in Hong Kong Waters (2018-19): Final Report. Hong Kong Cetacean Research Project Report to the Agriculture, Fisheries and Conservation Department, (2024).

[39] K.E. Klymus, C.M. Merkes, M.J. Allison, C.S. Goldberg, C.C. Helbing, M.E. Hunter, C.A. Jackson, R.F. Lance, A.M. Mangan, E.M. Monroe, A.J. Piaggio, J.P. Stokdyk, C.C. Wilson, C.A. Richter, Reporting the limits of detection and quantification for environmental DNA assays, Environmental DNA, 2 (2020) 271–282.

[40] Y. Zong, G. Huang, A.D. Switzer, F. Yu, W.W.S. Yim, An evolutionary model for the Holocene formation of the Pearl River delta, China, HOLOCENE, 19 (2009) 129–142.

[41] S. Oka, M. Miya, T. Sado, Gravity filtration of environmental DNA: A simple, fast, and power-free method, METHODSX, 9 (2022) 101838.

[42] T. Fukuzawa, H. Shirakura, N. Nishizawa, H. Nagata, Y. Kameda, H. Doi, Environmental DNA extraction method from water for a high and consistent DNA yield, Environmental DNA, 5 (2023) 627–633.

[43] M. Ushio, S. Furukawa, H. Murakami, R. Masuda, A.J. Nagano, An efficient early-pooling protocol for environmental DNA metabarcoding, Environmental DNA, 4 (2022) 1212–1228.

[44] M. Martin, CUTADAPT removes adapter sequences from high-throughput sequencing reads, EMBnet.journal, 17 (2011).

[45] S. Chen, Y. Zhou, Y. Chen, J. Gu, fastp: an ultra-fast all-in-one FASTQ preprocessor, BIOINFORMATICS, 34 (2018) i884–i890.

[46] B.J. Callahan, P.J. McMurdie, M.J. Rosen, A.W. Han, A.J.A. Johnson, S.P. Holmes, DADA2: High-resolution sample inference from Illumina amplicon data, NAT METHODS, 13 (2016) 581–583.

[47] R Core Team, R: A language and environment for statistical computing [Computer software], (2023).

[48] E. S. Wright, Using DECIPHER v2.0 to Analyze Big Biological Sequence Data in R, The R Journal, 8 (2016) 352.

[49] A.S. Tanabe, H. Toju, Two New Computational Methods for Universal DNA Barcoding: A Benchmark Using Barcode Sequences of Bacteria, Archaea, Animals, Fungi, and Land Plants, PLOS ONE, (2013).

[50] H. Wickham, ggplot2: elegant graphics for data analysis, Springer, New York, 2009.

[51] Z. Xia, A. Zhan, M.L. Johansson, E. DeRoy, G.D. Haffner, H.J. MacIsaac, Screening marker sensitivity: Optimizing eDNA-based rare species detection, DIVERS DISTRIB, 27 (2021) 1981–1988.

[52] V.S. Langlois, M.J. Allison, L.C. Bergman, T.A. To, C.C. Helbing, The need for robust qPCR-based eDNA detection assays in environmental monitoring and species inventories, Environmental DNA, 3 (2021) 519–527.

[53] T. Riaz, W. Shehzad, A. Viari, F. Pompanon, P. Taberlet, E. Coissac, ecoPrimers: inference of new DNA barcode markers from whole genome sequence analysis, NUCLEIC ACIDS RES, 39 (2011) e145–e145.

[54] M. Miya, Y. Sato, T. Fukunaga, T. Sado, J.Y. Poulsen, K. Sato, T. Minamoto, S. Yamamoto, H. Yamanaka, H. Araki, M. Kondoh, W. Iwasaki, MiFish, a set of universal PCR primers for metabarcoding environmental DNA from fishes: detection of more than 230 subtropical marine species, ROY SOC OPEN SCI, 2 (2015) 150088.

[55] E. Valsecchi, J. Bylemans, S.J. Goodman, R. Lombardi, I. Carr, L. Castellano, A. Galimberti, P. Galli, Novel universal primers for metabarcoding environmental DNA surveys of marine mammals and other marine vertebrates, Environmental DNA, 2 (2020) e72.

[56] C.J. Closek, J.A. Santora, H.A. Starks, I.D. Schroeder, E.A. Andruszkiewicz, K.M. Sakuma, S.J. Bograd, E.L. Hazen, J.C. Field, A.B. Boehm, Marine Vertebrate Biodiversity and Distribution Within the Central California Current Using Environmental DNA (eDNA) Metabarcoding and Ecosystem Surveys, FRONT MAR SCI, Volume 6 - 2019 (2019).

[57] S. Deng, X. Zhang, M. Liu, B. Lin, Y. Zhou, M. Liu, W. Lin, M. Lin, L. Dong, H. Kang, B. Liu, S. Chen, M. Ouyang, S. Jiang, J. Li, S. Li, Distribution pattern of cetaceans in the northern South China Sea based on visual surveys and environmental DNA metabarcoding, CONSERV BIOL, n/a (2025) e70060.

[58] G. Jeunen, M.D. Lamare, M. Knapp, H.G. Spencer, H.R. Taylor, M. Stat, M. Bunce, N.J. Gemmell, Water stratification in the marine biome restricts vertical environmental DNA (eDNA) signal dispersal, Environmental DNA, 2 (2020) 99–111.

[59] C.M. How, J.C. Ip, D. Deconinck, M. Zhao, M. Yan, J. Cheng, K.M.Y. Leung, L.L. Chan, J. Qiu, Refining Sampling Efforts for Fish Diversity Assessment in Subtropical Urban Estuarine and Oceanic Waters Using Environmental DNA With Multiple Primers, Environmental DNA, 6 (2024) e70013.

[60] Z. Yu, S. Ito, M.K. Wong, S. Yoshizawa, J. Inoue, S. Itoh, R. Yukami, K. Ishikawa, C. Guo, M. Ijichi, S. Hyodo, Comparison of species-specific qPCR and metabarcoding methods to detect small pelagic fish distribution from open ocean environmental DNA, PLOS ONE, 17 (2022) e0273670.

[61] C. Joseph, M.E. Faiq, Z. Li, G. Chen, Persistence and degradation dynamics of eDNA affected by environmental factors in aquatic ecosystems, HYDROBIOLOGIA, 849 (2022) 4119–4133.

[62] R.A. Collins, O.S. Wangensteen, E.J. O Gorman, S. Mariani, D.W. Sims, M.J. Genner, Persistence of environmental DNA in marine systems, COMMUN BIOL, 1 (2018) 185.

[63] P. Suarez-Bregua, M. Álvarez-González, K.M. Parsons, J. Rotllant, G.J. Pierce, C. Saavedra, Environmental DNA (eDNA) for monitoring marine mammals: Challenges and opportunities, FRONT MAR SCI, 9 (2022).

[64] G. Guri, J.L. Ray, A.O. Shelton, R.P. Kelly, K. Præbel, E. Andruszkiewicz Allan, N. Yoccoz, T. Johansen, O.S. Wangensteen, T. Hanebrekke, J. Westgaard, Quantifying the Detection Sensitivity and Precision of qPCR and ddPCR Mechanisms for eDNA Samples, ECOL EVOL, 14 (2024) e70678.

[65] X. Han, J. Chen, L. Wu, G. Zhang, X. Fan, T. Yan, L. Zhu, Y. Guan, L. Zhou, T. Hou, X. Xue, X. Li, M. Wang, H. Xing, X. Xiong, Z. Wang, Species distribution modeling combined with environmental DNA analysis to explore distribution of invasive alien mosquitofish (Gambusia affinis) in China, ENVIRON SCI POLLUT R, 31 (2024) 25978–25990.

[66] M. Riaz, M. Kuemmerlen, C. Wittwer, B. Cocchiararo, I. Khaliq, M. Pfenninger, C. Nowak, Combining environmental DNA and species distribution modeling to evaluate reintroduction success of a freshwater fish, ECOL APPL, 30 (2020).

[67] Q. Wu, J. Zhou, T. Komoto, T. Ishikawa, N. Goto, M.K. Sakata, D. Kitazawa, T. Minamoto, Opposite trends in environmental DNA distributions of two freshwater species under climate change, ECOSPHERE, 14 (2023).

[68] T.D. Lambert, S.P. Ellner, SDM meets eDNA: optimal sampling of environmental DNA to estimate species– environment relationships in stream networks, ECOGRAPHY, (2025).

[69] E. Pasanisi, D.S. Pace, A. Orasi, M. Vitale, A. Arcangeli, A global systematic review of species distribution modelling approaches for cetaceans and sea turtles, ECOL INFORM, 82 (2024) 102700.

