## Supplementary material for "Detecting the Indo-Pacific finless porpoises (*Neophocaena phocaenoides*) in Hong Kong waters using environmental DNA": Supplemental materials_1202.docx

**Contents**:

**Figure S1**. The structure of eDNA metabarcoding library constructed by the “early pooling” method.

**Figure S2**. Standard curve, the limit of detection (LOD) and limit of quantification (LOQ) of the primers and probe specific to *Neophocaena phocaenoides*.

**Figure S3**. Environmental parameters of the sampling locations of our survey area.

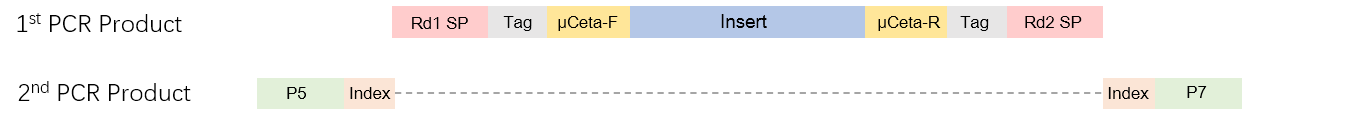

**Fig. S1.** **The structure of eDNA metabarcoding library for the “early pooling” method [1]**.

Tag sequences were appended in the first-round PCR, and the index sequences were appended in the second-round PCR. Both sequences were 8-bp sequences to differentiate samples. The detailed information is available in Ushio et al. (2022) [1].

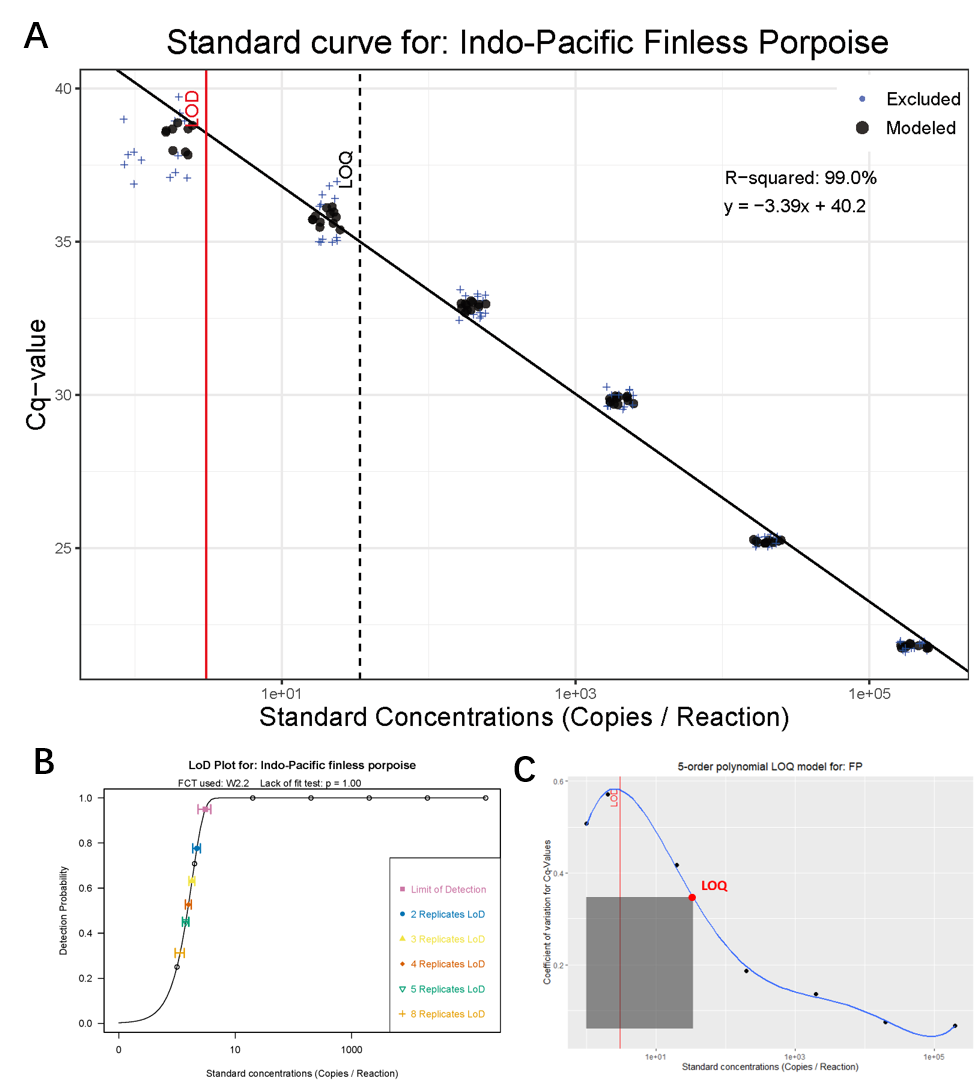

**Fig. S2**. **Standard curve, the limit of detection (LOD), and limit of quantification (LOQ) of the primers and probe specific to *Neophocaena phocaenoides*.** (**A**) Standard curve. The *x*-axis represents copy numbers of the target region per qPCR reaction, and the *y*-axis represents C_t_ value. Filled black points indicate data used to draw the linear regression line (50% of 24 replicates), while blue “+” symbols indicate those not used for the regression. Red solid line and black dashed line indicate LOD and LOQ, respectively. (**B**) Determination of LOD. Weibull type II model was selected as the best model. Different colors and symbols indicate LOD with different numbers of technical replicates. (**C**) Determination of LOQ. The x-axis represents copy numbers of the target region per qPCR reaction and the *y*-axis represents the coefficient of variation (C.V.). The blue curve indicates a 5th-order polynomial model (the best model), which was chosen based on the fitting residuals among 1-6th polynomial models (see Klymus et al. 2020 for details [2]). Red vertical line indicates LOD, and the red point indicates 35% C.V., of which copy number is defined as LOQ.

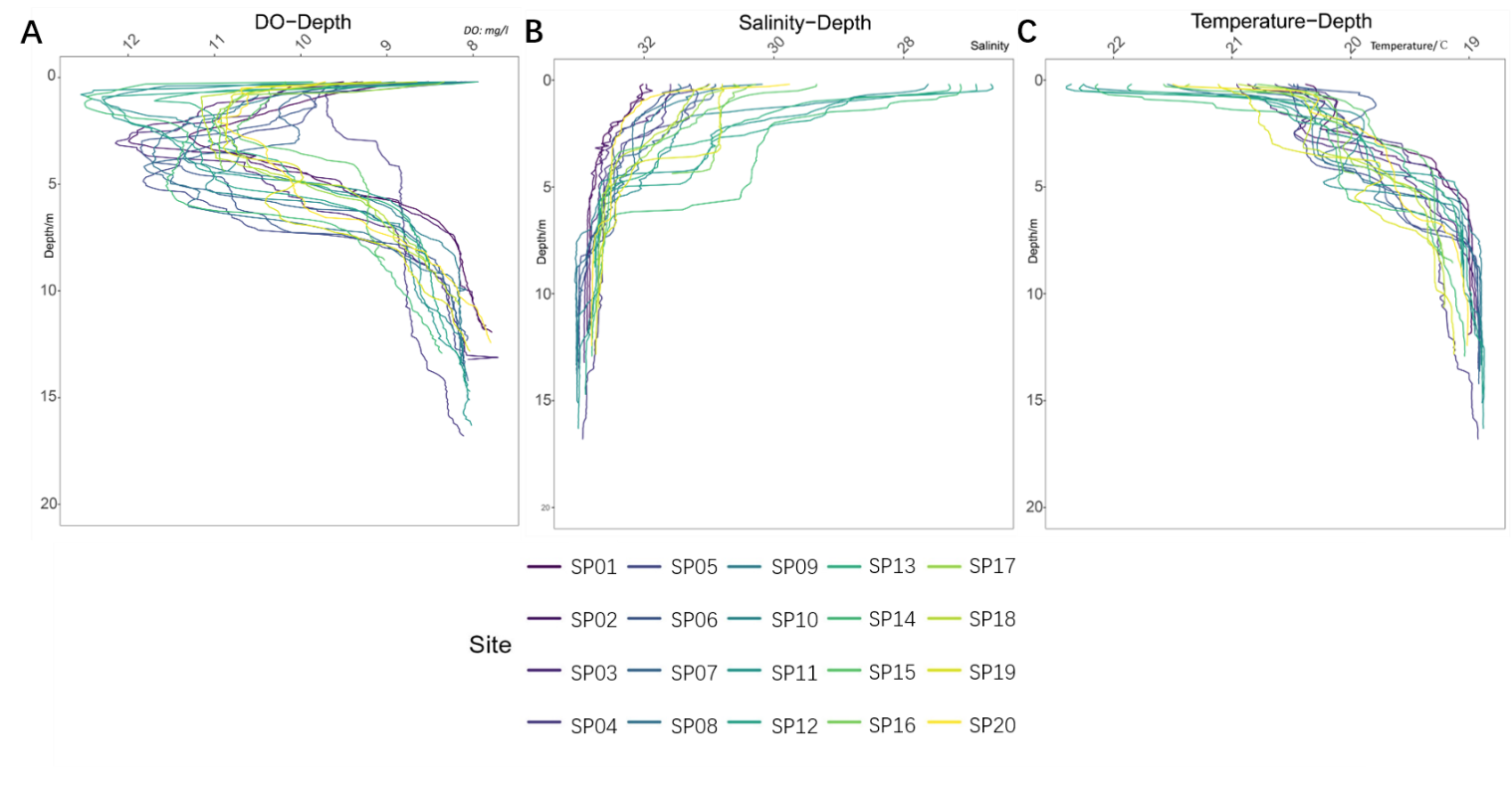

**Fig. S3**. **Environmental parameters of the sampling locations of our survey area.**

(**A**) Dissolved Oxygen (DO; mg/L), (**B**) Salinity (‰), (**C**) Water temperature (°C). Different colors indicate the different sampling locations. The *y*-axis indicates the water depth, while the *x*-axis indicates the environmental parameters.

References:

[1] M. Ushio, S. Furukawa, H. Murakami, R. Masuda, A.J. Nagano, An efficient early-pooling protocol for environmental DNA metabarcoding, Environmental DNA, 4 (2022) 1212-1228.

[2] K.E. Klymus, C.M. Merkes, M.J. Allison, C.S. Goldberg, C.C. Helbing, M.E. Hunter, C.A. Jackson, R.F. Lance, A.M. Mangan, E.M. Monroe, A.J. Piaggio, J.P. Stokdyk, C.C. Wilson, C.A. Richter, Reporting the limits of detection and quantification for environmental DNA assays, Environmental DNA, 2 (2020) 271-282.
